# Investigating Task-Free Functional Connectivity Patterns in Newborns Using functional Near-Infrared Spectroscopy

**DOI:** 10.1101/2023.09.05.555980

**Authors:** Homa Vahidi, Alexandra Kowalczyk, Kevin Stubbs, Melab Musabi, Sriya Roychaudhuri, Michaela Kent, Soume Bhattacharya, Sandrine de Ribaupierre, Keith St. Lawrence, Yalda Mohsenzadeh, Emma Duerden

## Abstract

**Significance:** Resting-state networks (RSN), particularly the sensorimotor network, begin to develop in the third trimester of pregnancy and mature extensively by term age. The integrity and structure of these networks have been linked to neurological health outcomes in neonates, highlighting the significance of monitoring RSN development. To this end, functional near-infrared spectroscopy (fNIRS) has emerged as a neuroimaging technique that utilizes near-infrared light to indirectly measure neural activity by detecting changes in oxygenated (HbO) and deoxygenated (HbR) hemoglobin concentrations. Compared to other imaging methods, fNIRS is non-invasive and allows for naturalistic monitoring of neural activity at the bedside, particularly in awake infants.

**Aim:** Use fNIRS to expand on previous findings regarding the development of functional networks in awake neonates.

**Approach:** fNIRS was acquired in 41 term-born neonates (17 females, gestational age range=36+0 to 42+1 weeks) within the first 48 hours after birth.

**Results:** Group level analysis of functional connectivity showed strong positive connectivity in most channel-pairs over the sensorimotor network, especially the left hemisphere (q < 0.05). Next, we examined the relationship between functional connectivity, gestational age and postnatal age, while controlling for sex and subject effects. Both gestational and postnatal age were found to be positively associated with an increase in functional connectivity in the sensorimotor RSN, especially in channels covering the posterior portion.

**Conclusions:** Our findings emphasize the importance of considering developmental changes in functional networks in awake infants. Moreover, our study demonstrates the potential of fNIRS as a valuable tool for studying neural activity in naturalistic settings in neonates.

## 1 Introduction

Task-free, or resting-state functional networks, which reflect the synchronized activity of different brain regions during rest, have been extensively studied in adult populations^1^ and are known to play a critical role in cognitive, motor, and sensory processes^2^. These networks emerge *in utero* and maintain a well-characterized structure throughout adulthood^3,4,5,6,7^. Alterations in these networks have been linked to various neurodevelopmental and psychiatric disorders^8,9^.

However, there continues to be a need for better characterizing these networks, especially in very early stages of development.

The first few hours and days of life mark a highly influential transition period for neonates. During this time, newborns are exposed to a multitude of novel sensory and motor experiences, laying the foundation for their cognitive and behavioral development^10^. Newborns receiving critical care soon after birth, particularly preterm babies (<37 weeks’ gestation), often lack the typical sensory and motor stimulations that healthy infants experience in the first few days of life^11^. This delay in sensory and motor enrichment may have implications for neurological and cognitive development. As such, characterizing resting-state functional networks in healthy newborns in this critical period is a first step toward gaining insights into early neurodevelopment.

Most of our current knowledge about resting-state networks comes from previous functional magnetic resonance imaging (fMRI) studies. Yet, conducting fMRI on newborns presents unique challenges, such as infants’ susceptibility to motion and sensitivity to loud noises. To overcome these limitations, functional near-infrared spectroscopy (fNIRS) emerges as a promising alternative that can expand our understanding of the development of resting-state networks, with the goal of monitoring these networks in a clinically relevant manner, particularly in neonates at risk of poor neurodevelopmental outcomes.

fNIRS allows for studies of cortical hemodynamic brain activity analogous to fMRI but in a more practical and comfortable manner, making it suitable for use in newborns even in clinical settings^12^. With higher temporal resolution than fMRI, fNIRS enables the determination of changes in the concentrations of both oxy-(HbO) and deoxy-hemoglobin (HbR), providing valuable insights into neuronal activity. Additionally, fNIRS can be employed for long-term recordings^13^, making it highly applicable in clinical settings for continuous monitoring of brain function in vulnerable neonates.

While a few studies have investigated resting-state networks in newborns using fNIRS^14,15,16^, less is known about the early development of these networks in the first few hours of life which is a critical period when the brain is most vulnerable to injury^17^. Uchitel and colleagues (2023) used high-density diffuse optical tomography (HD-DOT) to examine sleep states in relation to resting-state networks (RSN), offering valuable insights into the intricate interactions between neural activity and sleep processes in newborns and further indicate that bedside fNIRS is highly feasible in newborns. The sensorimotor network is expected to play a fundamental role in early sensorimotor development. More specifically, studying the development of the sensorimotor network in newborns is of particular interest, as this period marks a critical time for early motor exploration and sensory experiences that shape neural connections. By investigating the associations between gestational and postnatal age and sensorimotor connectivity, we hope to gain valuable insights into neurodevelopmental processes specific to this critical period. Given the importance of sensorimotor network development in the newborn brain, we aimed to use fNIRS to characterize this network. In a heterogeneous cohort of newborns who were born in a tertiary care center, we collected fNIRS data within the first 48 hours of life. We hypothesized that both gestational and postnatal age would be significantly associated with increased fNIRS-based connectivity in the sensorimotor network in these infants.

## 2 Methods

### 2.1 Study Setting and Participants

Participants were recruited from the Post-Partum Care Unit (PPCU) at Victoria Hospital, London, Ontario. Participants eligible for the study were healthy neonates born on or after 36 weeks of gestation. Neonates were excluded from the study based on the following criteria: congenital malformation or syndrome, antenatal exposure to illicit drugs, postnatal infection, and suspected brain injury. This study was approved by the Health Sciences Research Ethics Board of Western University and was conducted in accordance with the Declaration of Helsinki. All families provided written informed consent prior to data collection.

### 2.2 Demographic Data

Demographic and clinical information on maternal and newborn health were extracted from medical charts by a pediatric nurse or pediatrician. These data included: gestational age, postnatal age (age since birth), sex, and head size.

### 2.3 fNIRS Data Collection

Upon entering the PPCU, the healthcare team identified any families whose newborn would be eligible to participate in our study. All eligible families were first approached by their primary nurse to give verbal consent to be approached by researchers. After receiving written informed consent, the newborns’ head circumferences were measured and fit with an optode-prepopulated properly-sized fNIRS cap (34-38 cm). When possible, we recorded data after infants were fed to decrease the likelihood of motion and general fussiness.

We recorded task-free fNIRS signal from 61 infants at the bedside using a multichannel NIRSport2 system (NIRStar Software v14.0, NIRx Medical Technologies LLC, Berlin, Germany) at a sampling rate of 10.17 Hz. Our optode setup included 8 LED sources (760 and 850 nm) and 8 detectors which yielded 20 channels (10 per hemisphere). As confirmed with devfOLD^18^, our montage covered frontal and parietal regions of the sensorimotor network in both hemispheres. Based on devfOLD’s Lobar atlas at 60% specificity, 6 channels were located above the frontal lobe, 12 channels were located above the parietal portion, and 2 channels were included in both (See Fig. 1). Data were recorded for a minimum of 6 minutes and maximum of 10 minutes in each newborn to ensure stable and accurate functional connectivity calculations^19^. Data from 4 participants were excluded from preprocessing and analysis due to system malfunction and suboptimal calibration at the time of data collection. The remaining 57 (93%) infants comprised of 25 female and 32 male newborns with a mean gestational age of 39.01 ± 1.21 weeks and a mean postnatal age of 23.66 ± 11.75 hours.

**Figure 1.**
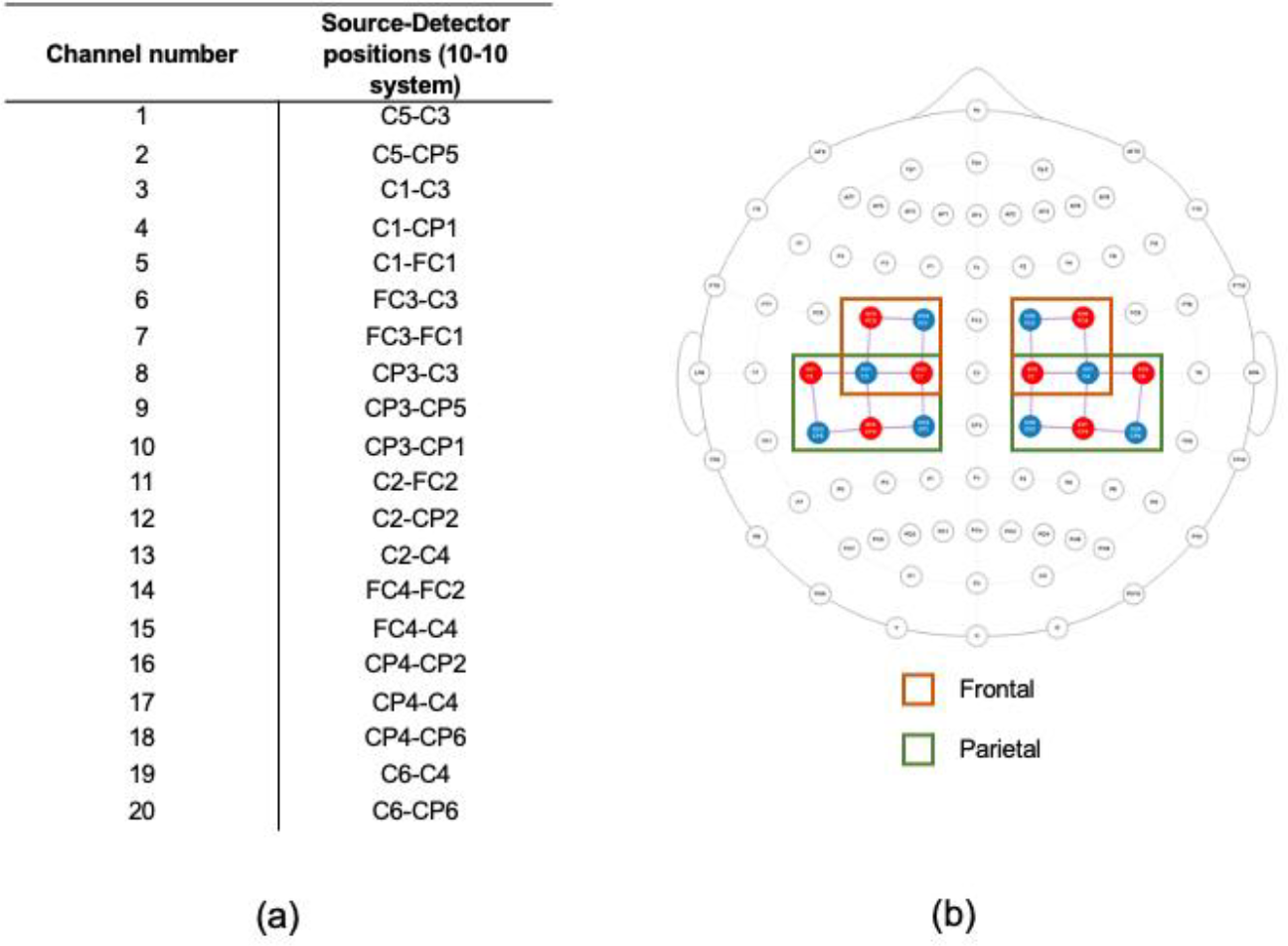
(a) 10-10 locations of source-detector pairs. (b) a 2-D top view of the montage used. Sources are shown in red, detectors are shown in blue, and channels between them are shown in purple. Channels overlaying the frontal portions of the sensorimotor network are highlighted by the orange rectangle and channels overlaying the parietal portions are highlighted by the green rectangle

### 2.4 fNIRS Preprocessing and Quality Assurance

All data pruning, preprocessing, and analysis were performed with MATLAB 2022b (The MathWorks Inc., Natick, Massachusetts, USA) using AnalyzIR Toolbox^20^, Homer2^21^, and in-house scripts.

#### 2.4.1 Channel/participant screening and exclusion

Raw data were first transformed to optical density. All channels in the 57 datasets were then screened for the presence of cardiac pulsation using an in-house cardiac detection method. This method combines scalp coupling index^22^, an indicator of how well each optode was contacting the scalp, with patterns in the frequency domain to flag channels without cardiac pulsation. For each dataset, all channels without cardiac pulsation were excluded. This process was verified through visual inspection. We fully excluded 12 datasets where no channels had detectable cardiac pulsation. See Figure. S1 for a group spatial map of all remaining channels. Next, for each of the remaining 45 datasets, we calculated scalp coupling index (SCI) and peak spectral power (PSP) in 5-s windows with 4-s overlaps for a cardiac range of 60–210 beats per minute, the cardiac range commonly reported in infants^23^. For each dataset, we used a SCI threshold of 0.1 and a PSP threshold of 0.03 to identify motion-free segments of at least 50-s. Brief motion artifacts (less than 2-s) were ignored during segmenting. While the SCI and PSP thresholds used were considerably lower than the defaults introduced in QT-NIRS^24^, we evaluated several different values (by visually inspecting each dataset before and after segmentation) and selected the most appropriate thresholds for our sample. Datasets were excluded if they did not produce any >50-s motion-free segments or if they had a total segment length less than 2.5-min, the minimum recommended dataset length for fNIRS resting-state functional connectivity analysis in infants^19^. This led to the removal of 4 datasets, leaving 41 datasets for further preprocessing and analysis.

The aforementioned steps led to a total of 16 (28%) datasets being excluded from further analyses. The remaining sample (n=41, 17 females) had a mean gestational age of 39.08 ± 1.31 weeks and a postnatal age of 22.93 ± 11.20 hours. We performed a chi-square test of independence to identify any sex differences between our dataset before and after participant exclusions. We additionally performed two 2-sample t-tests to identify any significant differences in gestational and postnatal ages. We found no statistically significant differences between the original dataset and the post-exclusion dataset.

#### 2.4.2 Preprocessing pipeline

The motion-free segments were preprocessed independently. Wavelet filtering with a standard deviation threshold of 0.8 (equivalent to a Homer iqr of 0.6) was applied to remove short motion spikes and slow drifts. Next, optical density data were transformed to estimated changes in HbO and HbR concentrations using the Modified Beer-Lambert Law. Age-appropriate partial pathlength factors were used (0.1063 for 760 nm and 0.0845 for 850 nm)^25^. Additionally, channel lengths were scaled to the cap size used. A bandpass filter (0.01 - 0.08 Hz) was applied to HbO and HbR separately. The data were then converted to estimated change in total hemoglobin concentration (HbT = HbO + HbR), which has been shown to have improved functional connectivity reproducibility across participants^26^. For each dataset, we normalized (mean = 0, SD = 1) and combined all segments. The final data were resampled to 4 Hz before calculating spontaneous functional connectivity (sFC). See Figure 2 for an overview of quality assurance and preprocessing steps.

**Figure 2.**
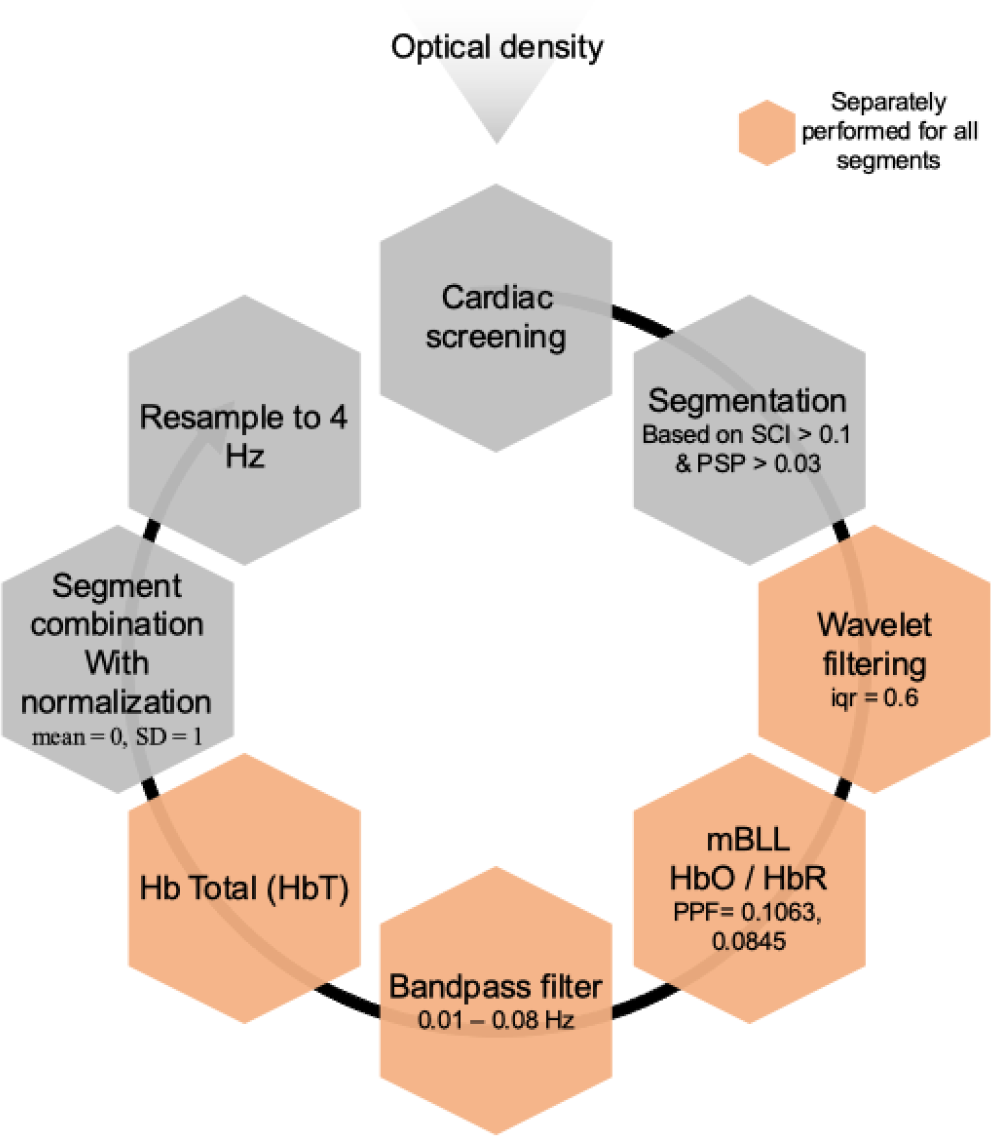
Overview of fNIRS data preprocessing steps. The black circular arrow indicates processing order starting with cardiac screening and ending with resampling to 4 Hz before sFC analysis.

### 2.5 Analysis Pipeline

#### 2.5.1 Spontaneous functional connectivity (sFC)

We used AnalyzIR Toolbox’s correlation method to calculate sFC. More specifically, for each dataset, we created a 20x20 symmetric correlation matrix by calculating Pearson’s R correlation coefficients between the time series of all possible channel-pairs. Correlation coefficients were then z-transformed. We performed a single-sample t-test for each unique channel-pair evaluating sFC > 0.

#### 2.5.2 Relating spontaneous functional connectivity (sFC) to gestational age and postnatal age

We used linear mixed effects (LME) modelling to evaluate the relationship between sFC and both gestational and postnatal age while accounting for variability due to biological sex and between-subject effects. Modelling was performed independently for each channel pair. Gestational age and postnatal age were included as fixed variables while subject ID and sex were included as random and grouping variables. Each channel-pair’s model produced a t-value and p-value for both gestational age and postnatal age, which indicates their linear relation to sFC across the sample.

#### 2.5.3 Statistical analysis

All statistical analyses were performed using MATLAB. We explored the relationship between sFC and both gestational age and postnatal age. These results were corrected for multiple comparisons using the false discovery rate (FDR) with a q-value set at 0.05. The number of channel-pairs (n=190) were used to calculate the degrees of freedom.

## 3 Results

### 3.1 Exploring Spatial Patterns in Spontaneous Functional Connectivity (sFC)

First, we examined the presence of any existing spatial sFC trends in our sample. We observed several significant positively correlated intra- and interhemispheric channels, especially in the left hemisphere (Figs. 3 and S2). Specifically, we observed positive connectivity (p<0.05) in 32/45 (∼71%) of all possible intrahemispheric channel-pairs in the left hemisphere, 20/45 (∼44%) of all possible intrahemispheric channel-pairs in the right hemisphere, and 27/100 (27%) of all possible interhemispheric channel-pairs. The sFC patterns in HbT were comparable to those seen in HbO (See Figs. S3 and S4). The sFC patterns in HbR were generally much weaker (See Figs. S5 and S6).

**Figure 3.**
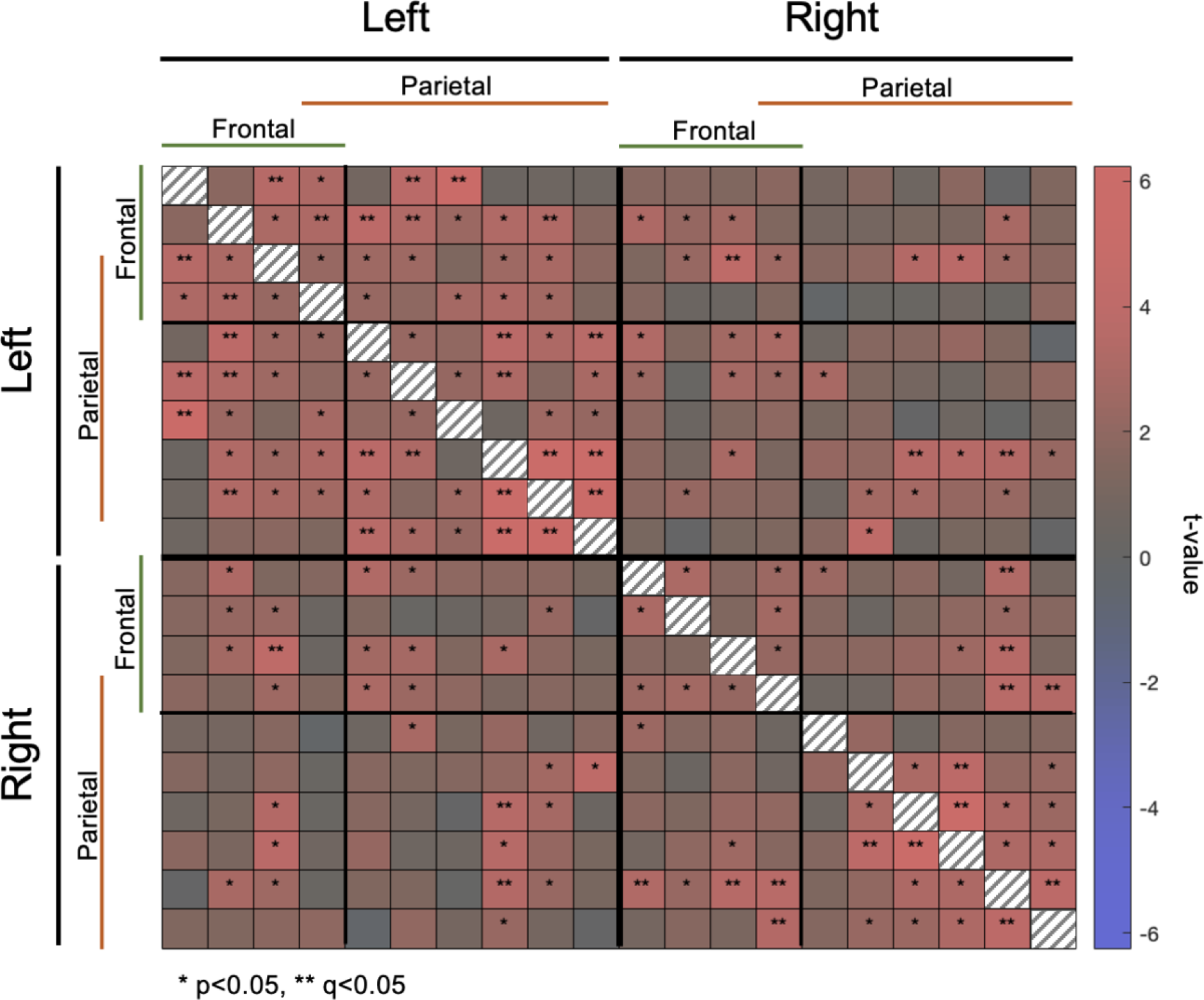
Matrix showing group spontaneous functional connectivity (sFC) for HbT (n=41). Channels overlaying the frontal and parietal portion of the sensorimotor network are denoted with the green and orange lines, respectively, with one channel overlaying both. False discovery rate (FDR) was used to correct for multiple comparisons. Channel-pairs that exhibited significant connectivity after FDR correction are highlighted using ** while channel-pairs with significant connectivity before FDR correction are highlighted using *. The color of matrix cells represents the t-value calculated for that channel-pair’s connectivity. The diagonal is not informative.

### 3.2 Relating Spontaneous Functional Connectivity (sFC) to Gestational Age and Postnatal Age

We identified several channel-pairs within and across the left and right hemispheres where increasing gestational age was significantly associated with both increasing and decreasing sFC (Figs. 4 and S7). Specifically, increasing gestational age was significantly associated with increasing sFC in 11/100 interhemispheric, 1/45 left intrahemispheric, and 1/45 right intrahemispheric channel-pairs. However, increasing gestational age was significantly associated with decreasing sFC in 8/100 interhemispheric, 5/45 left intrahemispheric, and 4/45 right intrahemispheric channel-pairs. The positive relationship between gestational age and sFC was mainly seen in channel-pairs connecting the parietal portion of the right sensorimotor network to the left sensorimotor network where 10 channel-pairs displayed this relationship. However, the negative relationship between gestational age and sFC was mainly seen in intrahemispheric channel pairs in the parietal portion of the left hemisphere as well as channel-pairs that connected the frontal portion of the right sensorimotor network to the parietal portion of the left hemisphere.

**Figure 4.**
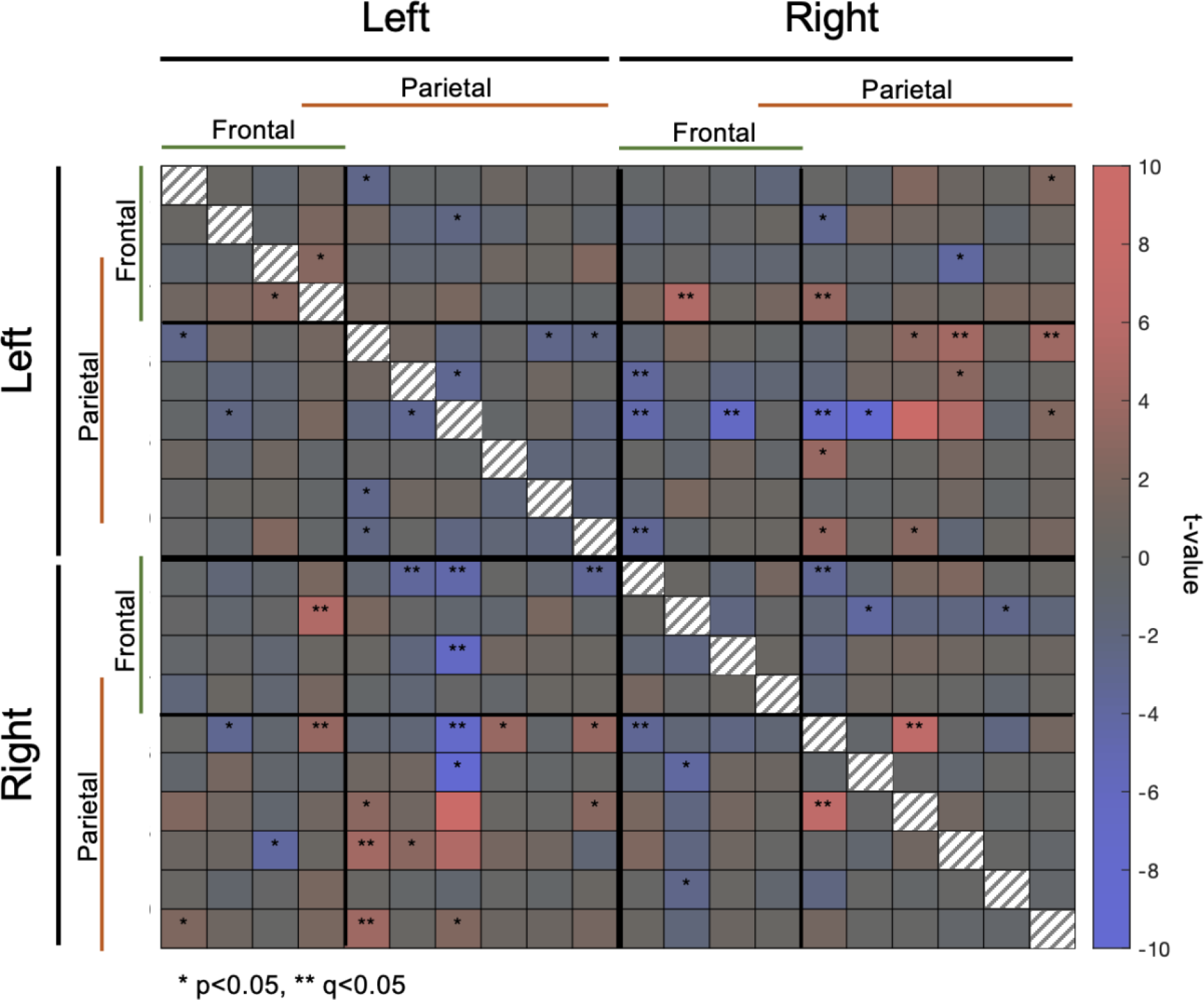
Matrix demonstrating gestational-age related patterns in spontaneous functional connectivity (sFC) (n=41). Channels overlaying the frontal and parietal portion of the sensorimotor network are denoted with the green and orange lines, respectively, with one channel overlaying both. False discovery rate (FDR) was used to correct for multiple comparisons. Channel-pairs whose connectivity exhibited a significant relationship with gestational age after FDR correction are highlighted using ** while channel-pairs showing a significant relationship before FDR correction are highlighted using *. The color of matrix cells represents the t-value calculated for that channel-pair. The diagonal is not informative.

We also identified several intrahemispheric and interhemispheric channel-pairs where increasing postnatal age was mainly associated with increasing sFC (Figs. 5 and S8). Specifically, increasing postnatal age was significantly associated with increasing sFC in 18/100 interhemispheric, 4/45 left intrahemispheric, and 5/45 right intrahemispheric channel-pairs. However, increasing postnatal age was significantly associated with decreasing sFC in 4/100 interhemispheric, 2/45 left intrahemispheric, and 2/45 right intrahemispheric channel pairs. The positive relationship between postnatal age and sFC was mainly seen in channel-pairs connecting the parietal portion of the right sensorimotor network to the left sensorimotor network and channel-pairs connecting the frontal portion of the right hemisphere to the parietal but not the frontal portion of the left hemisphere.

**Figure 5.**
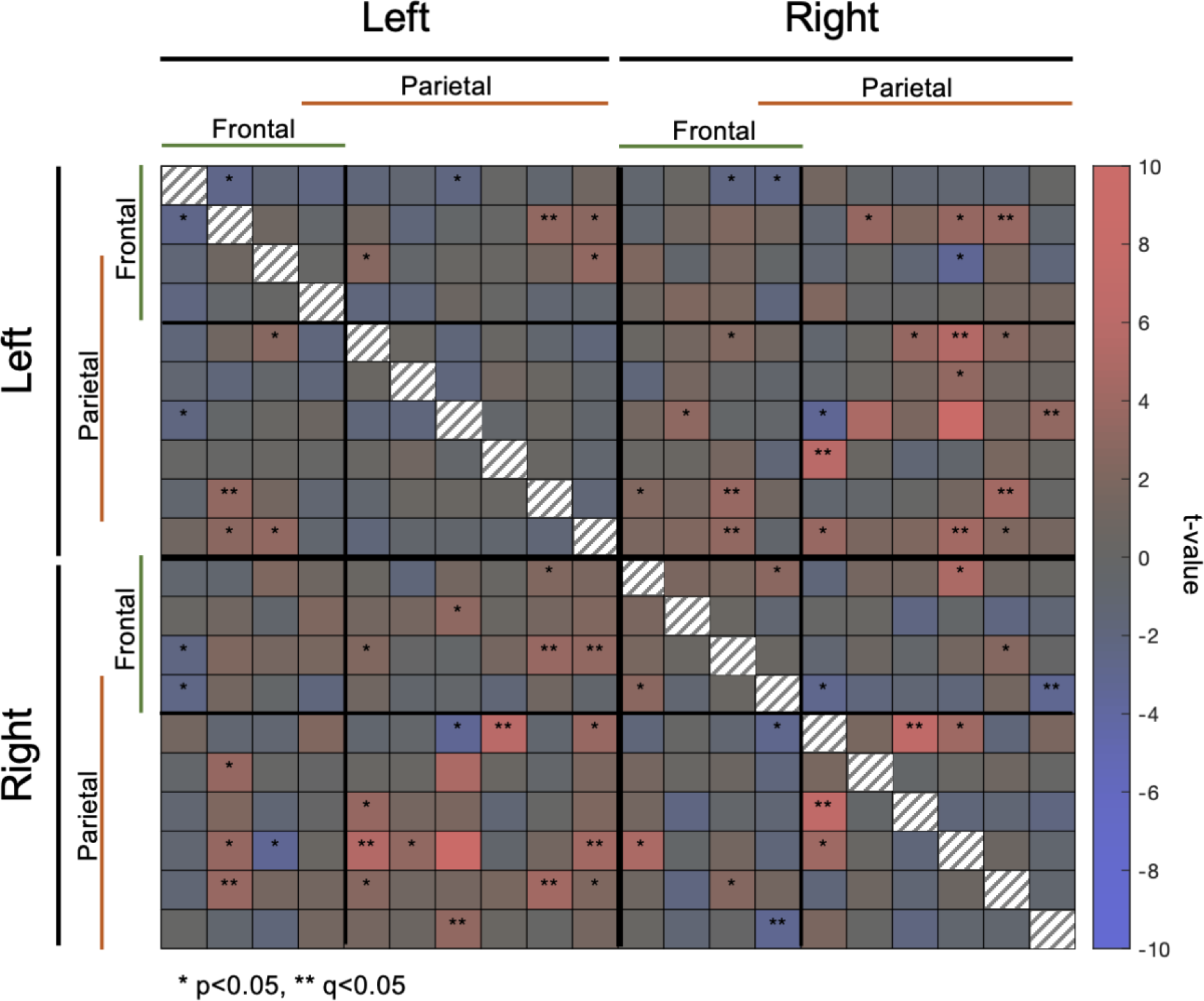
Matrix demonstrating postnatal-age related patterns in spontaneous functional connectivity (sFC) (n=41). Channels overlaying the frontal and parietal portion of the sensorimotor network are denoted with the green and orange lines, respectively, with one channel overlaying both. False discovery rate (FDR) was used to correct for multiple comparisons. Channel-pairs whose connectivity exhibited a significant relationship with postnatal age after FDR correction are highlighted using ** while channel-pairs showing a significant relationship before FDR correction are highlighted using *. The color of matrix cells represents the t-value calculated for that channel-pair. The diagonal is not informative.

## 4 Discussion

This work evaluated functional changes in sensorimotor RSN connectivity in the first few days of life in healthy newborns. We collected 10-minute fNIRS recordings from healthy newborns within the first 48 hours of life in the hospital setting. We evaluated the relationship between sFC and gestational and postnatal age, while accounting for variability due to biological sex and between-subject effects. We identified a strong, left-lateralized sensorimotor RSN for which connectivity, particularly in channels covering the parietal portion of the network, had a positive relationship with older gestational and postnatal ages. We also assessed sensorimotor RSN developmental changes during the first hours of life spent in the *ex-utero* environment. Bilateral sensorimotor RSN connectivity was significantly strengthened during the first days of life, pointing to the rapid development of the sensorimotor network and highlighting the importance of mapping out the trajectories of structural and functional development in this critical period of life. We mapped the connectivity between all channel-pairs in the sensorimotor RSN in our sample. Similar to previous fMRI and fNIRS studies in neonates^3,27,13^, we observed strong positive interhemispheric and intrahemispheric connectivity in the sensorimotor network. We observed the strongest connectivity between channel-pairs within the left hemisphere. Human sensorimotor functions are often left lateralized depending on handedness preference ^28^. As more than 90 percent of the general population prefers using their right hand to perform most motor tasks^29^, this lateralization may be determined early on in development. In fact, ultrasound monitoring of fetal arm movements has indicated a potential asymmetry in hand preference as early as the second trimester^30^. Additionally, this preference has been associated with a left-dominant lateralization in the sensorimotor cortex^31^. As such, the strong intrahemispheric connectivity we observed in the left hemisphere could be an early indicator of laterality in the sensorimotor network.

We further examined the relationship between gestational age, postnatal age and sFC in the sensorimotor network. Older gestational ages were associated with a modest increase in bilateral connectivity in the sensorimotor network. More specifically, we observed a positive relationship between gestational age and sFC in several channel-pairs that connected the parietal portion of right hemisphere to channels across the left hemisphere. The positive relationship observed between gestational age and connectivity was somewhat weaker than those reported in previous fMRI studies^3^. Unexpectedly, we also observed a negative relationship between gestational age and sFC in a few channel-pairs connecting the frontal portion of the right hemisphere to the left hemisphere. Our findings could be due to the older gestational age range of our sample compared to previous studies. More specifically, previous fMRI and fNIRS studies reporting a strong positive relationship between gestational age and sFC have also included preterm neonates and thus had a wider gestational age range^3,6^. In turn, the current study in healthy term-born infants might be capturing sFC in a narrower developmental window, with fewer maturational changes that could be evidenced with fNIRS^27^. Development of cortical RSNs is reflective of synchronous maturation of cortical gray matter and white matter. Essential to this process *in utero* is subplate connectivity reflected in synaptogenesis and thalamo-cortical projections. This period is marked by rapid synaptogenesis, which is offset by neuronal apoptosis and pruning of weaker synapses^32,33^. Additionally, a key factor in RSN development is myelination. Broadly, subcortical areas myelinate first followed by the posterior cortex and frontal cortex. More specifically, myelination begins in the central sulcus and extends toward the posterior cortex followed by the frontotemporal locations. The structural and functional maturation of these processes are shaped by neuronal activity and external stimuli in the extrauterine environment^32,34^.

Postnatal age (i.e., hours since birth) was associated with widespread unilateral and bilateral connectivity across channels covering the sensorimotor network. Findings may reflect heightened neuronal activity in the postnatal environment due to the exposure to novel environmental stimuli in the first hours of life. Ferradal and colleagues (2016), utilizing HD-DOT, recorded task-free oscillations in brain activity in newborns during a similar postnatal period (i.e., within the first 2 days of life) also reported strong interhemispheric connectivity in middle temporal, visual, and auditory RSNs^16^. However, limited intrahemispheric connectivity was reported for the same networks. While our findings were primarily localized to the territory of the sensorimotor network, we found increasing interhemispheric and some evidence for left-sided intrahemispheric connectivity at older postnatal ages in this RSN during natural sleep/rest. Intra-hemispheric connectivity may be a key factor supporting hemispheric specialization^35^. The first few days of life are characterized by jerky and non-goal directed general movements, in addition to early motor reflexes^36^. The first days and weeks of life are characterized by the development of coordinated motor movements^37,38^. Absence of these sensorimotor behaviors may be an indicator for poor neurodevelopment later in life^39^. In turn, the use of fNIRS at the bedside may identify early biomarkers for typical sensorimotor development.

### 4.1 Study Limitations

Our study included a heterogenous sample of day-old newborns who were tested with a standardized fNIRS protocol. Despite the challenges faced by recruiting this vulnerable population, our results provide evidence for the emergence of robust RSNs as well as inter- and intra-hemispheric connectivity that is associated with brain maturational stages. However, our study had several limitations that are inherent to fNIRS data collection in infants. Namely, we excluded datasets and/or channels of subpar quality that led to variable number of viable channels in each dataset. The regression method we employed to assess the relationship between sFC and both gestational and postnatal age accounted for the number of channel-pairs used.

Nonetheless, it is advisable for future studies to employ larger sample sizes to corroborate and expand upon our findings.

Our montage was confined to motor, premotor, and sensory cortical regions. Consequently, we were unable to examine developmental changes in other RSNs during this crucial phase. Prior fMRI studies involving newborns have showcased consistent and robust developmental pathways for networks encompassing sensory and motor regions (e.g., sensorimotor, auditory, and visual networks)^3^. However, the developmental trajectories identified for more complex networks (e.g., the Default Mode Network) are more variable and tend to be challenging to examine due to the coarse spatial resolution of fNIRS. As a result, for the scope of this study, we opted to concentrate solely on the sensorimotor network; however, future work could also include whole-head coverage to better characterize the sensorimotor networks in relation to other RSNs to better understand early cortical connectomics.

Further, newborns’ sensory and motor function was not assessed. All newborns enrolled in this study were examined by a pediatrician and were healthy in relation their motor development. Furthermore, subject-related effects were accounted for in our regression model. Incorporating standardized neurodevelopmental assessments could expand on how sensorimotor RSN characteristics predict motor outcomes.

### 4.2 Conclusions

This study aimed to advance our understanding of the development of sensorimotor RSNs in healthy newborns using fNIRS. By exploring functional connectivity within the sensorimotor RSNs and how it is associated with gestational and postnatal age, this study contributes to our understanding of normative brain development during early yet critical stages of life. While our study demonstrates the feasibility of using fNIRS at the bedside in neonates, our findings also highlight the challenges associated with fNIRS data collection and analysis in postpartum care centers. Given the utility of fNIRS in healthy newborns and neonates impacted by critical illness, our study highlights the need for improved fNIRS methodologies tailored for this vulnerable population.

## 6 Data and Code Availability

The data and code used in this study is available upon request.

## 8 Acknowledgements

The authors extend their sincerest gratitude to all the families and staff members of the post-partum unit who made this research possible. The authors would also like to acknowledge New Frontiers in Research Fund, NSERC CRSNG, Brain Canada Foundation, and CIHR whose financial contributions were essential to this study.

