## Supplemental Figures for "Investigating Task-Free Functional Connectivity Patterns in Newborns Using functional Near-Infrared Spectroscopy"

**Supplementary Materials**

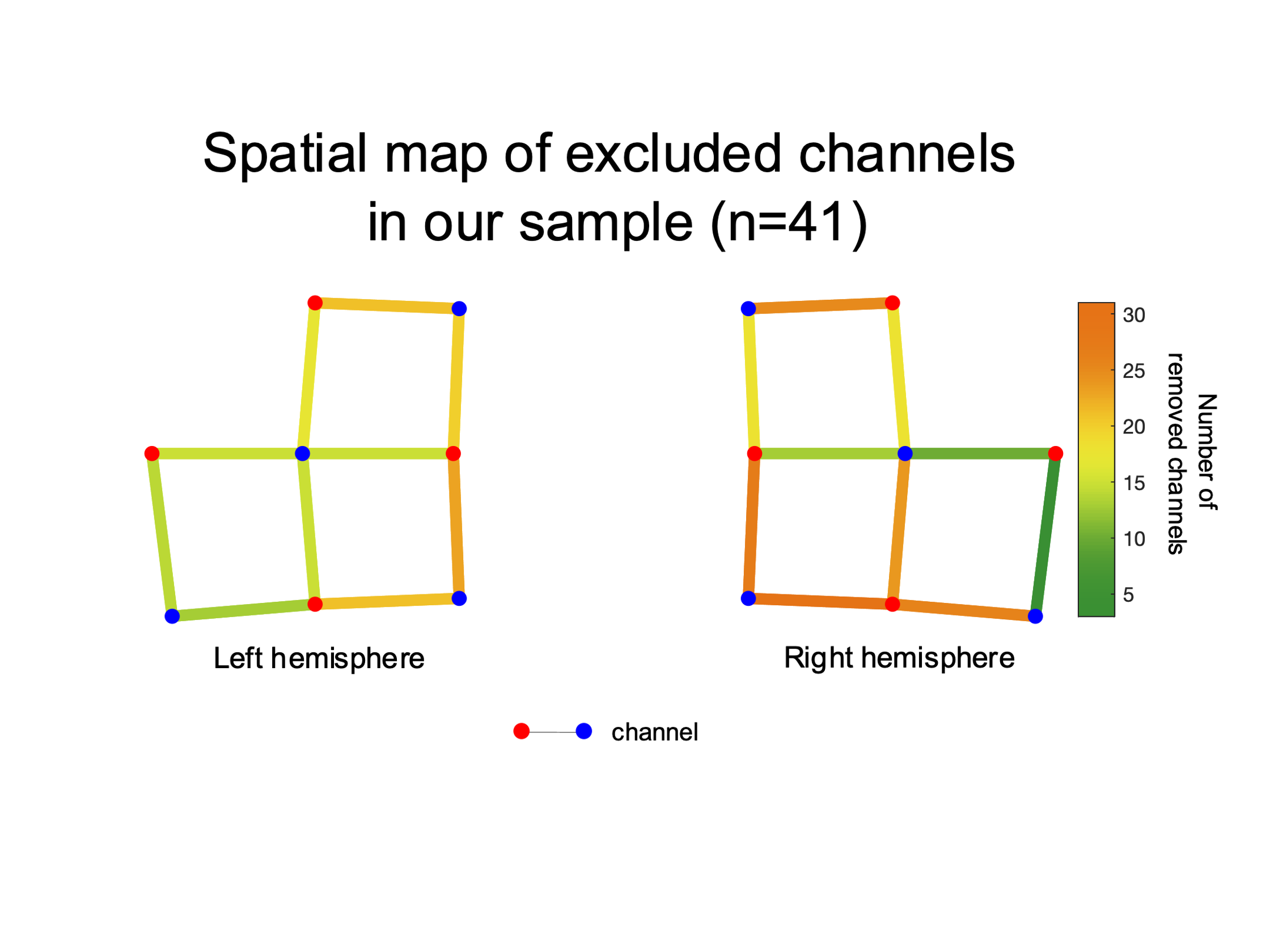

**Figure S1. Group spatial map of channel exclusions (n=41).** The color of the lines depicts the number of infants in which that channel was excluded. Channels depicted in green were preserved in most infants while channels depicted in orange were excluded in most infants.

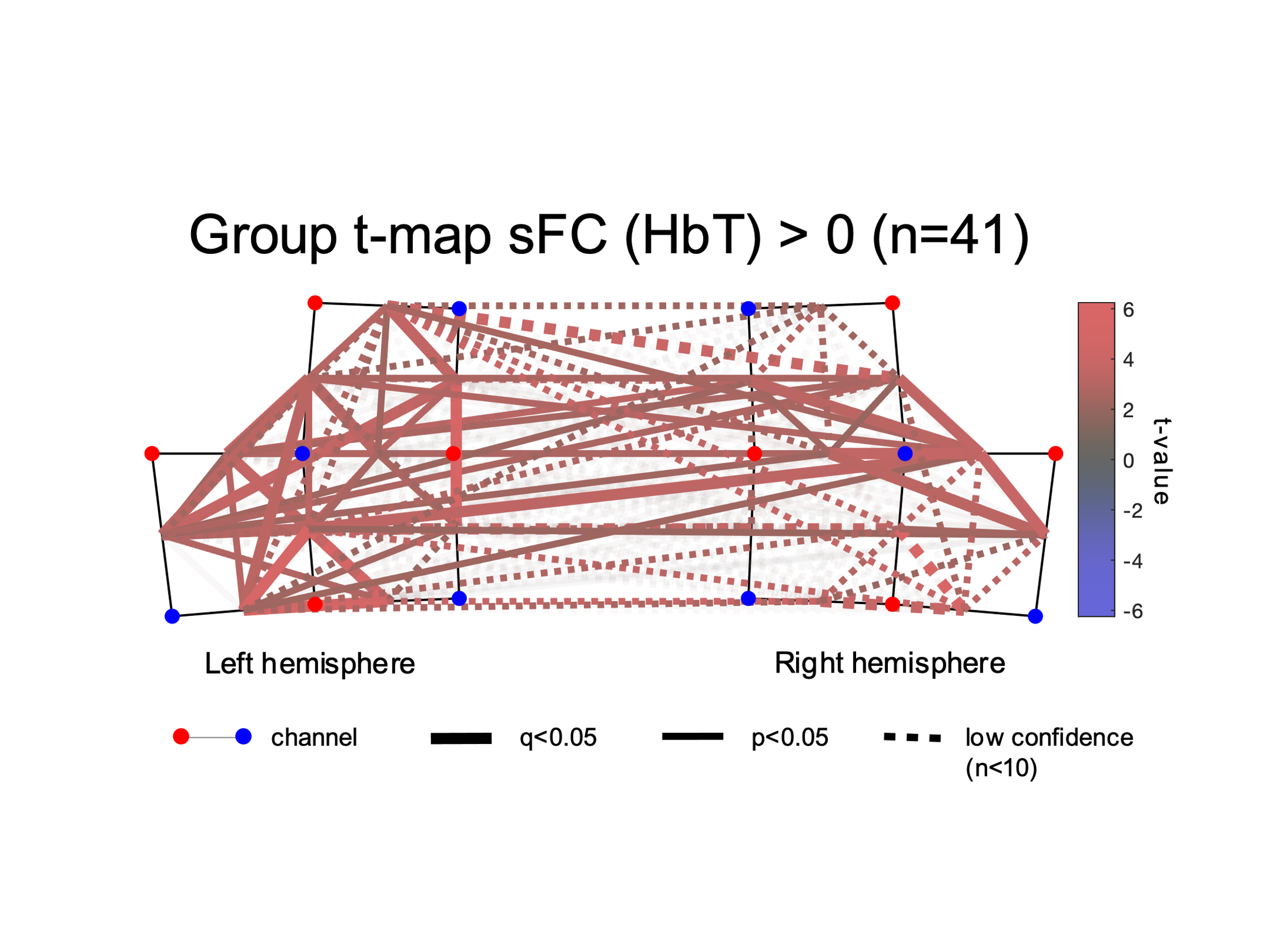

**Figure S2. Group t-map showing spontaneous functional connectivity (sFC) spatial patterns for HbT (n=41).** Channel-pairs displaying a significant positive or negative sFC are depicted in red and blue lines, respectively. The false discovery rate (FDR) was used to correct for multiple comparisons. Channel-pairs that exhibited significant connectivity after FDR correction are drawn as thick lines while channel-pairs with significant connectivity before FDR correction are denoted with thin lines. The color of the lines represents the t-value calculated for that channel-pair’s connectivity. Channel-pairs that had fewer than 10 datapoints are depicted with dotted lines, which signifies the small sample used for that specific channel pair. Channels pairs that were not significant have been faded to increase clarity.

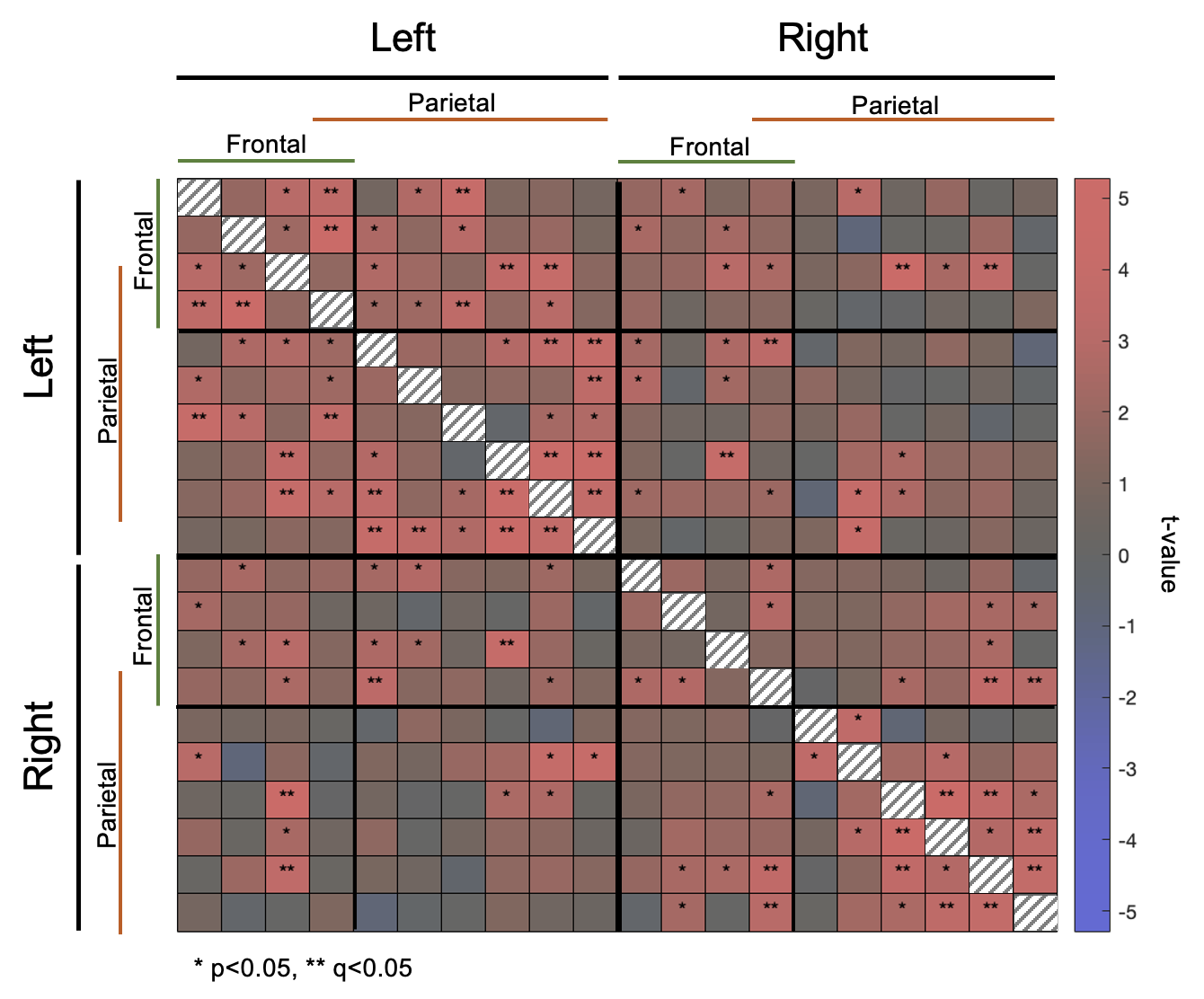

**Figure S3. Matrix showing group spontaneous functional connectivity (sFC) for HbO (n=41).** Channels overlaying the frontal and parietal portion of the sensorimotor network are denoted with the green and orange lines, respectively, with one channel overlaying both. False discovery rate (FDR) was used to correct for multiple comparisons. Channel-pairs that exhibited significant connectivity after FDR correction are highlighted using ** while channel-pairs with significant connectivity before FDR correction are highlighted using *. The color of matrix cells represents the t-value calculated for that channel-pair’s connectivity. The diagonal is not informative.

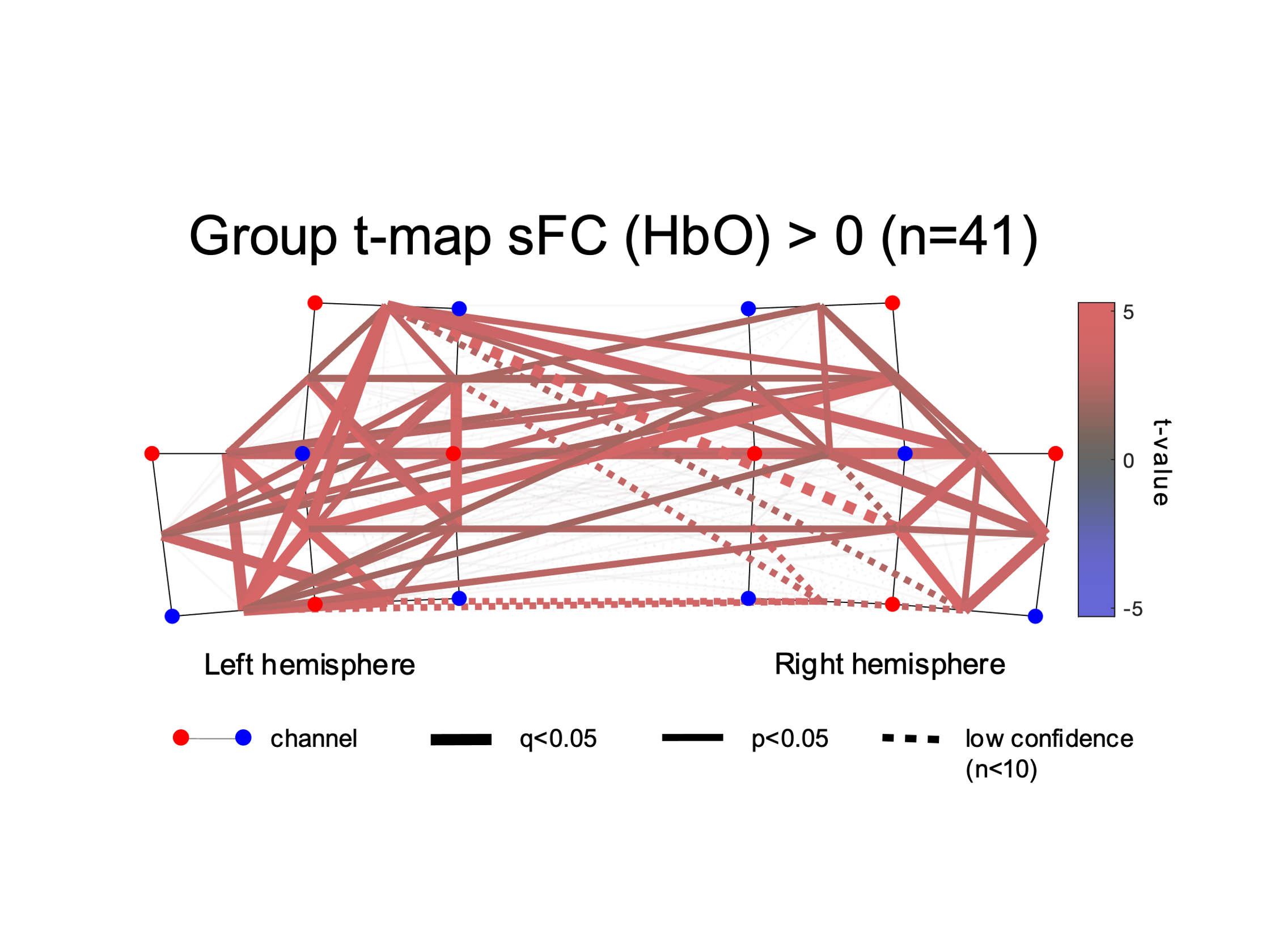

**Figure S4. Group t-map showing spontaneous functional connectivity (sFC) spatial patterns for HbO (n=41).** Channel-pairs displaying a significant positive or negative sFC are depicted in red and blue lines, respectively. The false discovery rate (FDR) was used to correct for multiple comparisons. Channel-pairs that exhibited significant connectivity after FDR correction are drawn as thick lines while channel-pairs with significant connectivity before FDR correction are denoted with thin lines. The color of the lines represents the t-value calculated for that channel-pair’s connectivity. Channel-pairs where one or both channels were missing (due to subpar quality) were not included in the analysis. Channel-pairs that had <10 datapoints are depicted with dotted lines, which signifies the small sample used for that specific channel pair. Channels pairs that were not significant have been faded out to increase clarity.

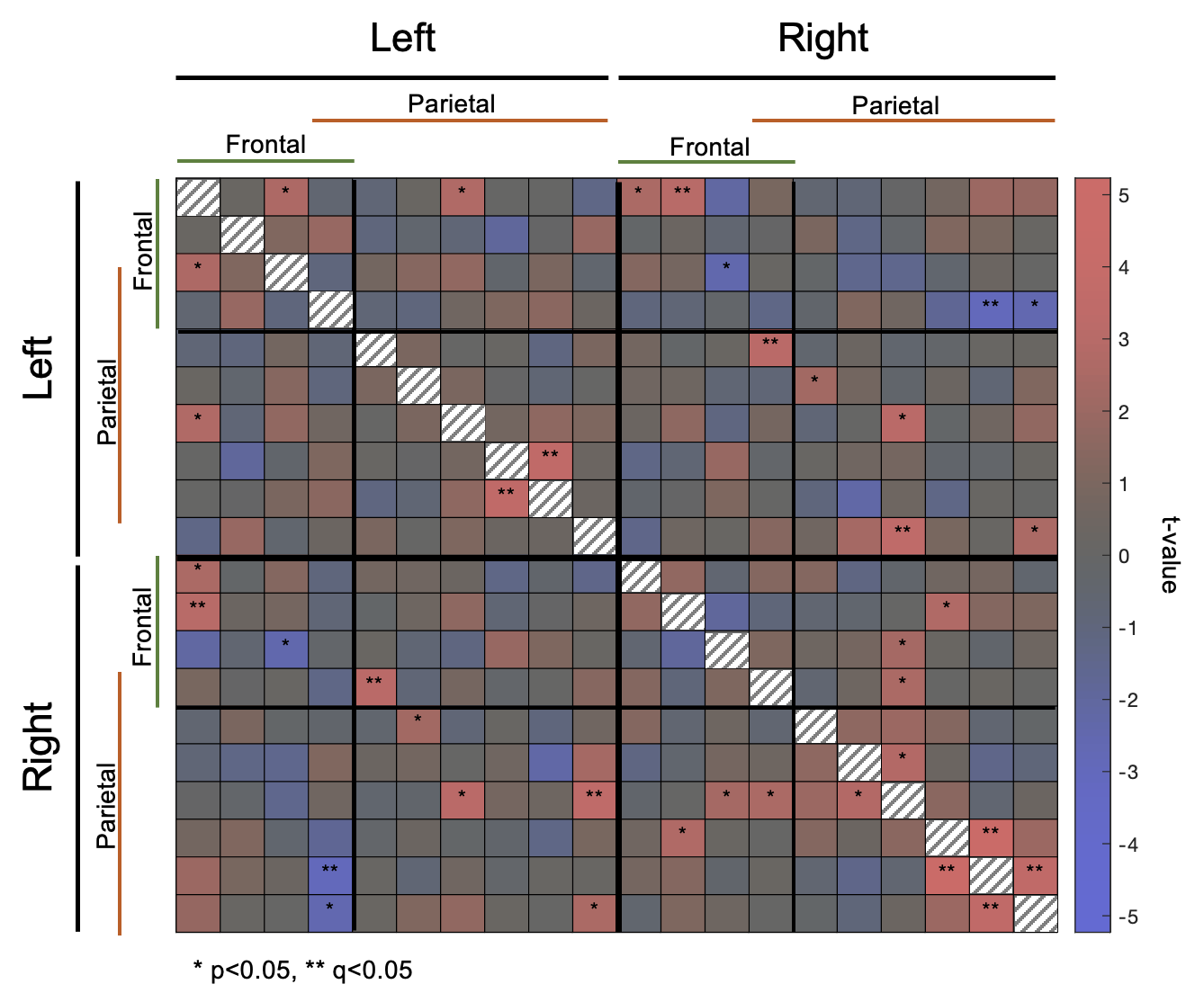

**Figure S5. Matrix showing group spontaneous functional connectivity (sFC) for HbR (n=41).** Channels overlaying the frontal and parietal portion of the sensorimotor network are denoted with the green and orange lines, respectively, with one channel overlaying both. False discovery rate (FDR) was used to correct for multiple comparisons. Channel-pairs that exhibited significant connectivity after FDR correction are highlighted using ** while channel-pairs with significant connectivity before FDR correction are highlighted using *. The color of matrix cells represents the t-value calculated for that channel-pair’s connectivity. The diagonal is not informative.

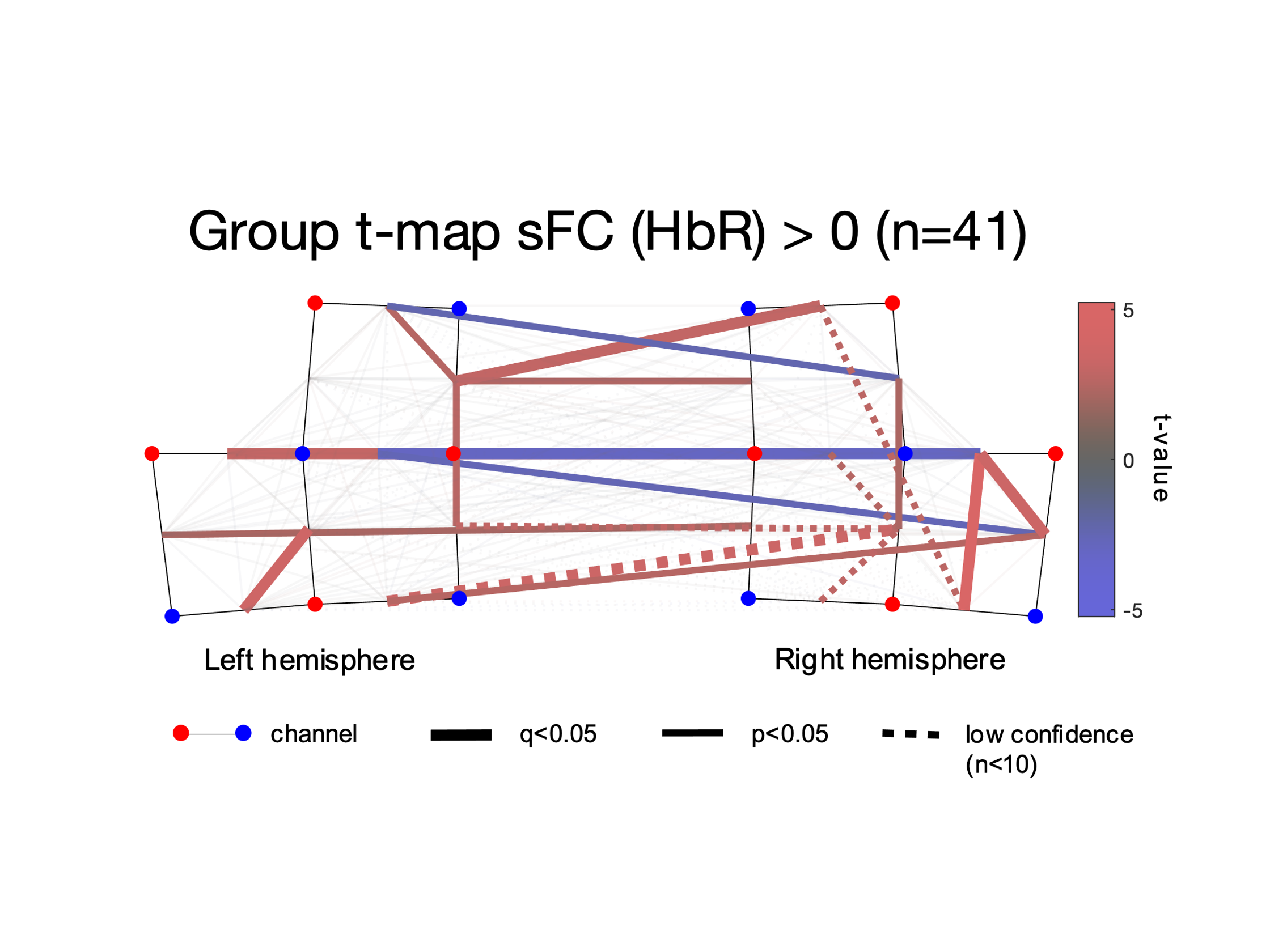

**Figure S6. Group t-map showing spontaneous functional connectivity (sFC) spatial patterns for HbR (n=41).** Channel-pairs displaying a significant positive or negative sFC are depicted in red and blue lines, respectively. The false discovery rate (FDR) was used to correct for multiple comparisons. Channel-pairs that exhibited significant connectivity after FDR correction are drawn as thick lines while channel-pairs with significant connectivity before FDR correction are denoted with thin lines. The color of the lines represents the t-value calculated for that channel-pair’s connectivity. Channel-pairs where one or both channels were missing (due to subpar quality) were not included in the analysis. Channel-pairs that had <10 datapoints are depicted with dotted lines, which signifies the small sample used for that specific channel pair. Channels pairs that were not significant have been faded out to increase clarity.

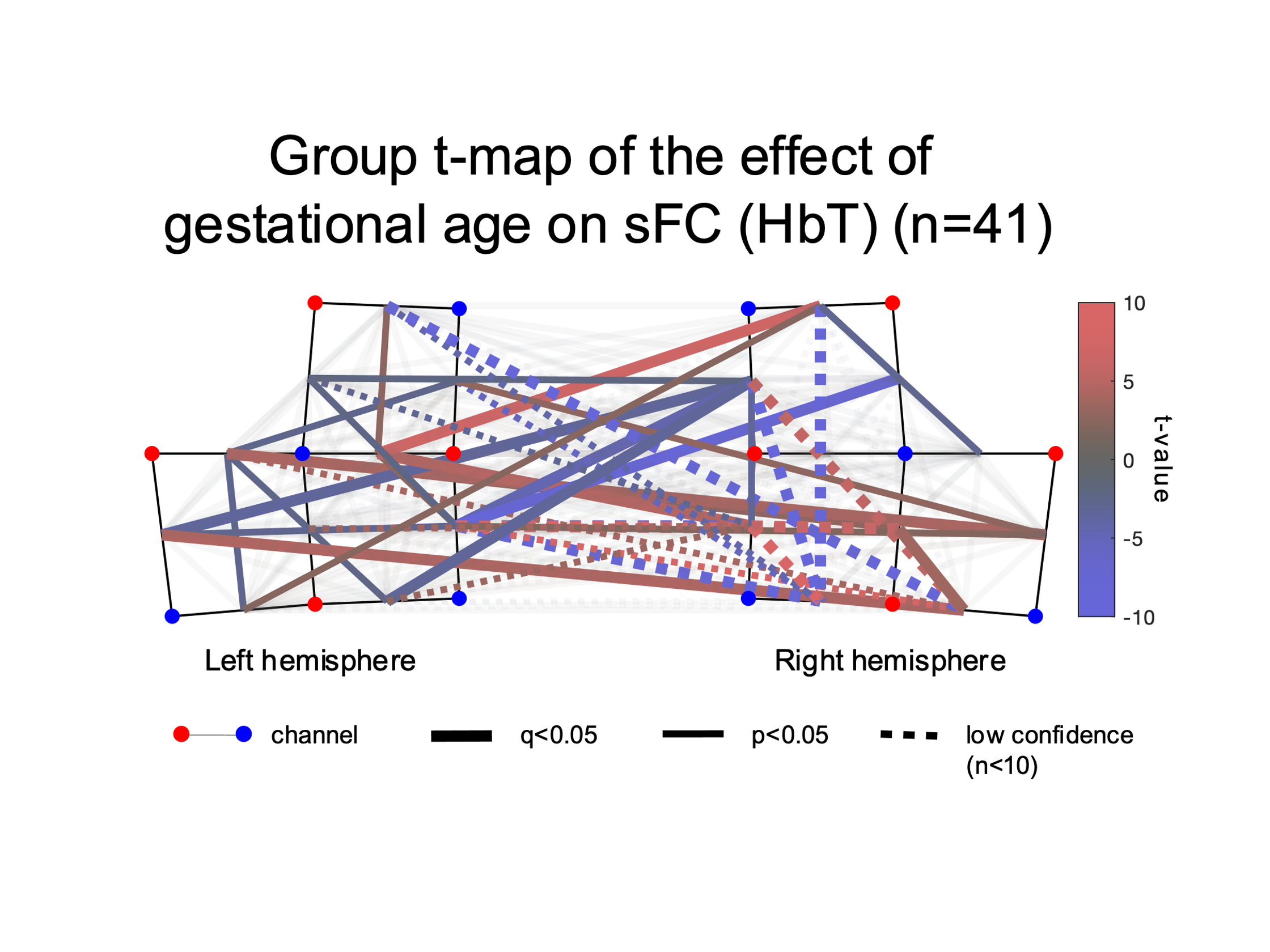

**Figure S7. Group regression analysis demonstrating gestational-age related patterns in spontaneous functional connectivity (sFC) (n=41).** Channel-pairs displaying a significant positive or negative effect of gestational age on sFC are depicted in red and blue lines, respectively. The false discovery rate (FDR) was used to correct for multiple comparisons. Channel-pairs that exhibited significant connectivity after FDR correction are drawn as thick lines while channel-pairs with significant connectivity before FDR correction are denoted with thin lines. The color of the lines represents the t-value calculated for that channel-pair’s connectivity. Channel-pairs that had fewer than 10 datapoints are depicted with dotted lines, which signifies the small sample used for that specific channel pair. Channels pairs that were not significant have been faded out to increase clarity.

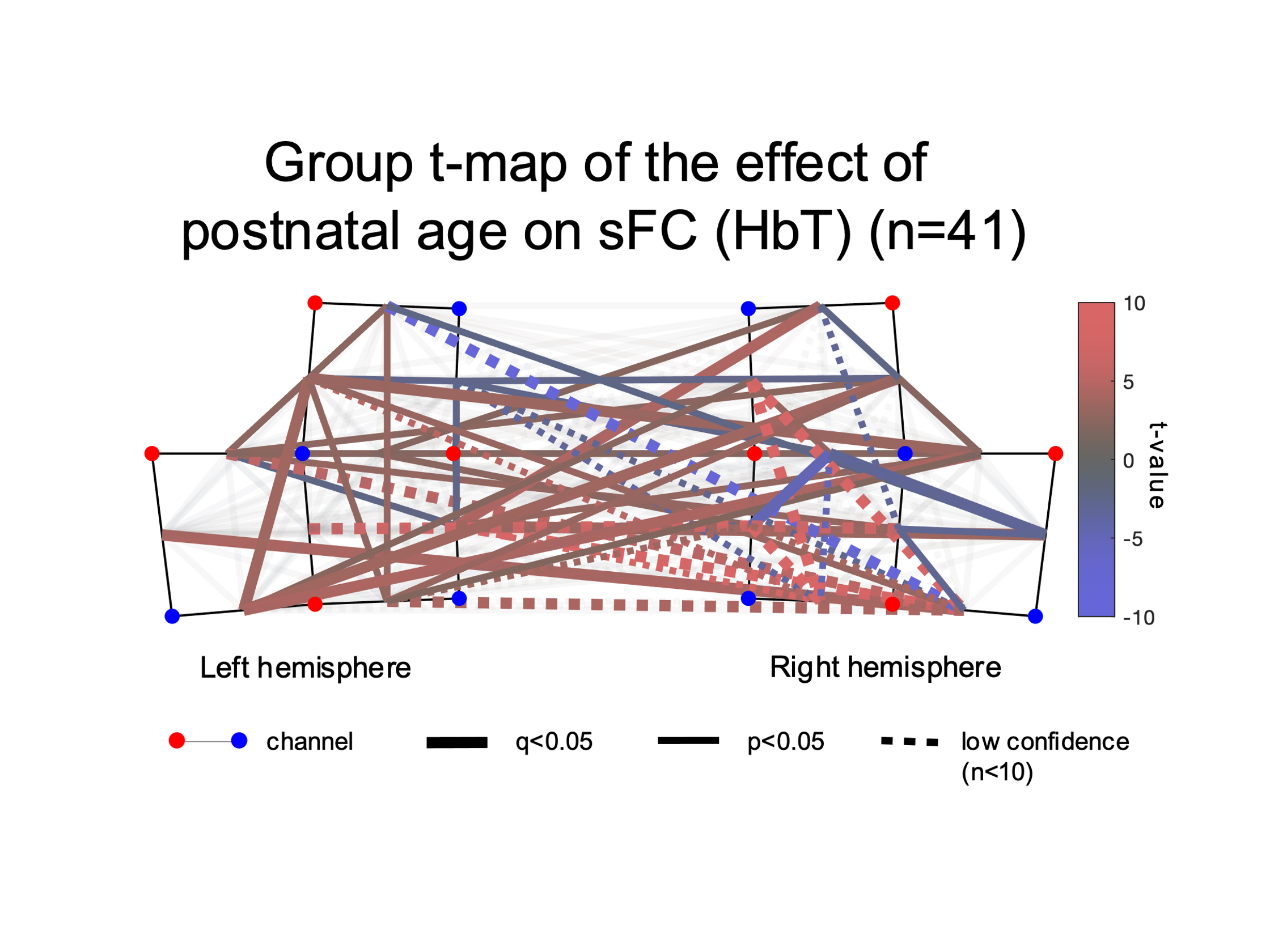

**Figure S8. Whole montage regression analysis demonstrating postnatal-age related patterns in functional connectivity (sFC) (n=41).** Channel-pairs displaying a significant positive or negative effect of postnatal age on sFC are depicted in red and blue lines, respectively. The false discovery rate (FDR) was used to correct for multiple comparisons. Channel-pairs that exhibited significant connectivity after FDR correction are drawn as thick lines while channel-pairs with significant connectivity before FDR correction are denoted with thin lines. The color of the lines represents the t-value calculated for that channel-pair’s connectivity. Channel-pairs that had fewer than 10 datapoints are depicted with dotted lines, which signifies the small sample used for that specific channel pair. Channels pairs that were not significant have been faded out to increase clarity.
